# The historical domestication of a *Clostridium botulinum* strain used for the industrial production of botulinum neurotoxin

**DOI:** 10.64898/2026.02.26.708219

**Authors:** Paul Keim, Roxanne Nottingham, Miriam A. Guevara, Ely F. Miller, Amy J. Vogler, Charles H. D. Williamson, Theresa Smith, Richard G. Posner, Sabine Pellett, Richard E. Lenski, Jason W. Sahl

## Abstract

Laboratory production of botulinum neurotoxin (BoNT) began more than 90 years ago for medical, military, and later pharmaceutical applications, creating one of the longest-running examples of microbial cultivation under sustained human control. A single *Clostridium botulinum* Group I lineage, Army Hall A (AHA), was established by the U.S. military in 1942 and later gave rise to Hall A-hyper (HAH), the strain widely used for pharmaceutical BoNT production. We analyzed more than 1,000 *C. botulinum* genomes to identify AHA’s closest relatives and to infer the most recent common ancestor (MRCA) shared with its nearest wild lineage. Relative to this MRCA, AHA accumulated nearly 100 genetic changes, including 46 single-nucleotide substitutions, 44 small insertions or deletions, and several large deletions and structural variants that led to the loss of more than 80 genes. A nonsense mutation in *mutS* generated a hypermutator phenotype that accelerated mutation rates and increased genetic diversity. This event occurred early in the lineage’s domestication and appears to have facilitated laboratory-adaptive traits, including the lack of sporulation and increased BoNT yield. Competition experiments under standard growth conditions confirmed substantial laboratory adaptation, with AHA exhibiting a strong fitness advantage over a close wild relative. Together, these results reconstruct the genomic trajectory of a bacterium evolving under prolonged human-mediated selection and provide a genome-resolved example of microbial domestication. The findings show how laboratory conditions, industrial selection, and changing mutation rates can jointly shape bacterial evolution, and they offer a general framework for understanding domestication and laboratory adaptation across microbial systems.

**Significance:** Human-mediated species domestication has been central to the development of civilization, with well-known examples in plants and animals. Microbes have also been domesticated, often unintentionally and before the advent of modern microbiology. The military and pharmaceutical applications of botulinum neurotoxin led to the sustained cultivation and eventual domestication of a high toxin-producing *Clostridium botulinum* strain that remains widely used today. We reconstruct the genomic changes along this trajectory and identify a key early mutation that generated a hypermutator phenotype, increasing genetic variation available for human-directed selection. These findings reveal fundamental evolutionary processes shaping bacterial genomes under long-term human control. This genome-resolved example of microbial domestication offers a general framework for understanding laboratory adaptation in both evolutionary and applied microbiology.

## Introduction

Domestication represents a powerful process for human-driven evolution, in which selection, both intentional and inadvertent, has reshaped the genomes, behaviors, and physiologies of diverse species (1, 2). Through sustained selection on heritable variation, humans have transformed wolves into numerous dog breeds (3), wild grasses into high-yield crops (4), and wild yeasts into efficient fermenters used in beer, wine, and bread production (5). Central to these processes is the availability of genetic diversity, which provides the substrate on which selection acts to produce populations specialized for human-constructed environments (2, 6). Although the genetic and evolutionary bases of animal and plant domestication are well characterized, the corresponding processes in microbes remain largely unexplored (7). Microbial domestication arises when laboratory or industrial procedures for propagation impose selective pressures that favor traits advantageous for growth, stability, or productivity under these conditions.

A striking example of microbial domestication is *Clostridium botulinum*, which produces a neurotoxin (BoNT) that has been repurposed from its natural role as a virulence factor to a manufactured biologic used for therapeutic and cosmetic applications (8). Historically, BoNT’s extreme potency motivated its use in both offensive and defensive biological-weapons programs, which inadvertently laid the foundation for its biomedical utility (9). The domesticated *C. botulinum* lineages that have been maintained for industrial BoNT production therefore provide an intriguing model for studying how human intervention, driven by both military and medical motivations, has shaped microbial adaptation and genome evolution in controlled environments.

Foodborne botulism is an intoxication illness frequently associated with improperly prepared foods that permit anaerobic growth of *C. botulinum* and the production of BoNT (10). As food-preservation efforts expanded during the early 20th century, so too did outbreaks of botulism (11). A notable 1919–1920 outbreak linked to canned olives stimulated systematic research to understand and mitigate this emerging threat. Among those engaged in this effort was Dr. Ivan C. Hall, who assembled an extensive collection of *C. botulinum* isolates (12). Several of these strains were shared with Dr. Leland McClung at Indiana University and tested by Dr. Elizabeth McCoy for toxin productivity. This led to the discovery of one isolate that exhibited particularly high toxin yields (13), which became known simply as “the Hall strain.” The strain was later obtained by Dr. J. H. Mueller (Harvard Medical School) and transferred in 1942 to the U.S. Army at Camp Detrick, Maryland, where it was used to generate culture supernatants containing high levels of toxin and the first biochemical isolation of botulinum neurotoxin type A (14), which was used for both offensive and defensive weapons research programs (15) including the development of antitoxins and toxoid vaccines (16, 17). By 1956, investigators had noted “Hall A’s” failure to sporulate, necessitating frequent subculturing to maintain the organism (17). These early reports thus document two phenotypic traits—high toxin yield and loss of sporulation—that remain central to the strain’s continued use in pharmaceutical BoNT production.

When the U.S. eliminated its biological weapons program in 1972, a subculture of the “Hall A” strain was transferred to the Food Research Institute at the University of Wisconsin-Madison, where further studies on the production, purification, and crystallization of this toxin were conducted (18, 19). This subculture was eventually re-named Hall A-hyper to reflect its abundant toxin yields (20). The pure crystalline toxin was used to produce the first crystal structure of a botulinum toxin (21), and a collaboration with Dr. Alan Scott enabled the first use of botulinum toxin as a therapeutic agent, known as Oculinum (22). That Hall A botulinum toxin batch was the first commercial use of therapeutic BoNT/A1, including the initial BOTOX® product (23). The Hall A-hyper strain was selectively distributed to private, public, and commercial entities, and its A1 toxin is likely the most common BoNT produced globally for both therapeutic and cosmetic uses (24).

In this study, we conducted a phylogenomic analysis to elucidate the evolutionary processes underlying the domestication of the Army Hall A (AHA) strain and its central role in both military and pharmaceutical BoNT production. Our analysis reveals a distinct phylogenetic branch defined by 46 single-nucleotide mutations, 44 small indels, a chromosomal translocation (25), and two large deletions. These mutations are consistent with both genome decay and adaptation accompanying a niche shift from nature to the laboratory and industrial environments. AHA domestication was accelerated soon after isolation by a nonsense mutation in the *mutS* gene, which produced a hypermutator phenotype that increased the rate of genome decay and facilitated the accumulation of adaptive mutations. Laboratory populations lack the influx of external genetic variants by horizontal gene transfer, and thus this hypermutator phenotype provided an internal source of diversity that accelerated adaptation in clonal lineages. The resulting strain exhibits loss of sporulation, enhanced BoNT production, and high fitness relative to a related wild-type strain under laboratory conditions.

## Results

### Evolution of Army Hall A from a wild ancestor

Understanding how the AHA strain evolved from a wild Group I *C. botulinum* (*C. parabotulinum)* ancestor requires identifying other closely related strains to define their most recent common ancestor (MRCA). AHA and its derived Hall A-hyper (HAH) strain are a small subset of the broad diversity within *Clostridium botulinum* Group I (*SI Appendix*, Fig. S1a) that has been shaped by gene transfer, mutation, and niche-specific selection (26). The evolutionary relationships among 20 closely related AHA relatives are defined by 135 SNPs (*SI Appendix,* Fig. S1b). One 12-member clade includes different accessions of the lab strain NCTC7272/ATCC19397 (11), while a second 4-member clade contains food-associated outbreak isolates, including BrDuraA from Argentina. The AHA genome is separated from these relatives along with two derived HAH accessions. These three Hall A strains are separated from their clade’s MRCA by a relatively long branch. The first AHA genome (CP000727) was sequenced >20 years ago, and we confirmed its accuracy by sequencing a subclone from the same Army cell bank (182-M2). The X58540 (GCA_016798345.1) genome is the closest known relative to AHA that was recently isolated from nature (27). While their core genomes are nearly identical (99.999%), the differences are critical for understanding the laboratory evolution of AHA from its wild ancestor.

### Chromosomal synteny between AHA and its closest relative

Fig. 1a shows the evolutionary relationships between AHA, X58540, ATCC19397, and BrDuraA based on SNPs in their core genomes. This phylogenetic model assigns individual point mutations to branches and enables mapping of additional genomic variation. We divided all genomic differences between AHA and X58540 into two strain-specific branches (Fig. 1a) by comparison to outgroups to define their MRCA. The AHA branch has 46 SNPs and 44 indels, while the X58540 branch has only 4 SNPs and no indels (Table 1). Chromosomal alignments show nearly complete synteny between X58540 and AHA with three exceptions (Fig. 1b), including a previously identified translocation (25). The translocation involves 16 genes moved by ∼200 kbp, with no gene loss (*SI Appendix,* Fig. S2). Although AHA retains all genes in this region, the translocation could still affect gene expression or regulation.

**Fig. 1.**
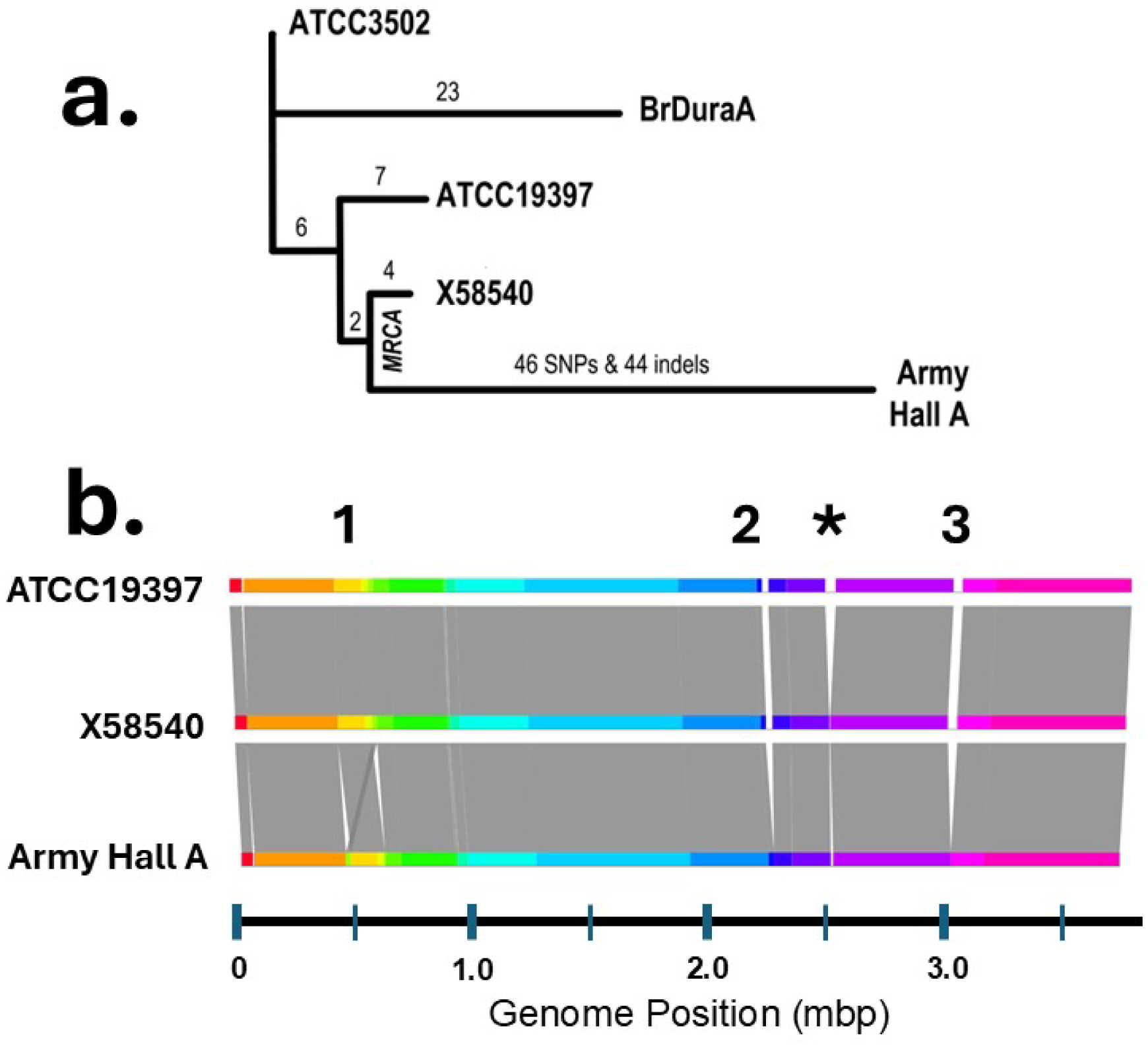
Genomic differences between Army Hall A and its close relatives. **a,** A phylogenetic diagram showing the relationship of the AHA genome to three near relatives: BrDuraA, ATCC19397, and X58540. ATCC3502 was used as an outgroup to assign ancestral versus derived SNPs. The phylogenetically informative SNPs in Table 1a correspond to those labeled here on different branches. Indels are indicated only for the AHA branch, but these and others are also shown in Table 1b. **b,** A full genome progressive Mauve alignment of the AHA, X58540, and ATCC19397 genomes illustrates their synteny and where the few structural differences exist. The #1 marks a 16-gene AHA-specific translocation previously reported by Fang et al. (25), while two large deletions in the AHA genome are labeled #2 (21 genes) and #3 (61 genes); they are detailed in Dataset S2 along with two other single-gene deletions. The asterisk (*) marks a multiple-gene deletion unique in ATCC19397. An HTML-powered graphic of the AHA-specific translocation region (MUMmer alignment) is shown in *SI Appendix,* Fig. S2.

**Table 1.**
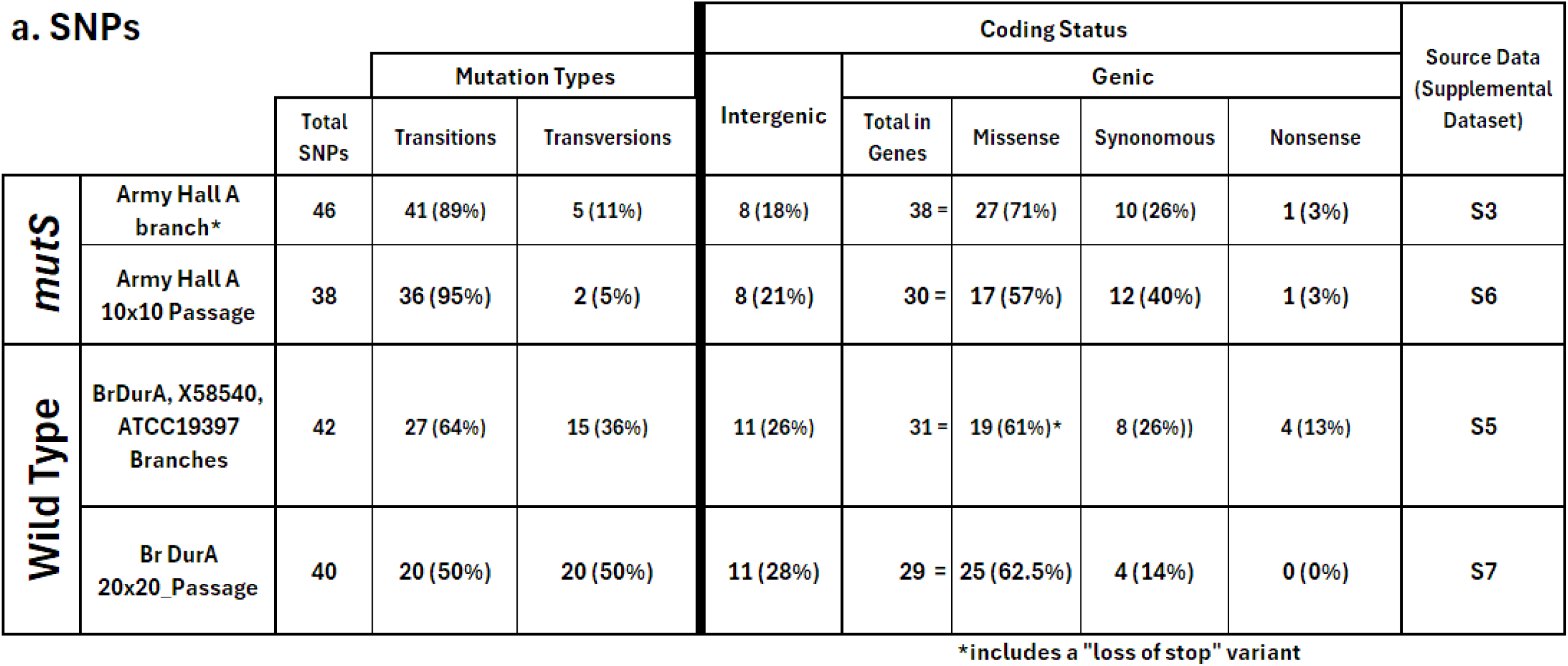
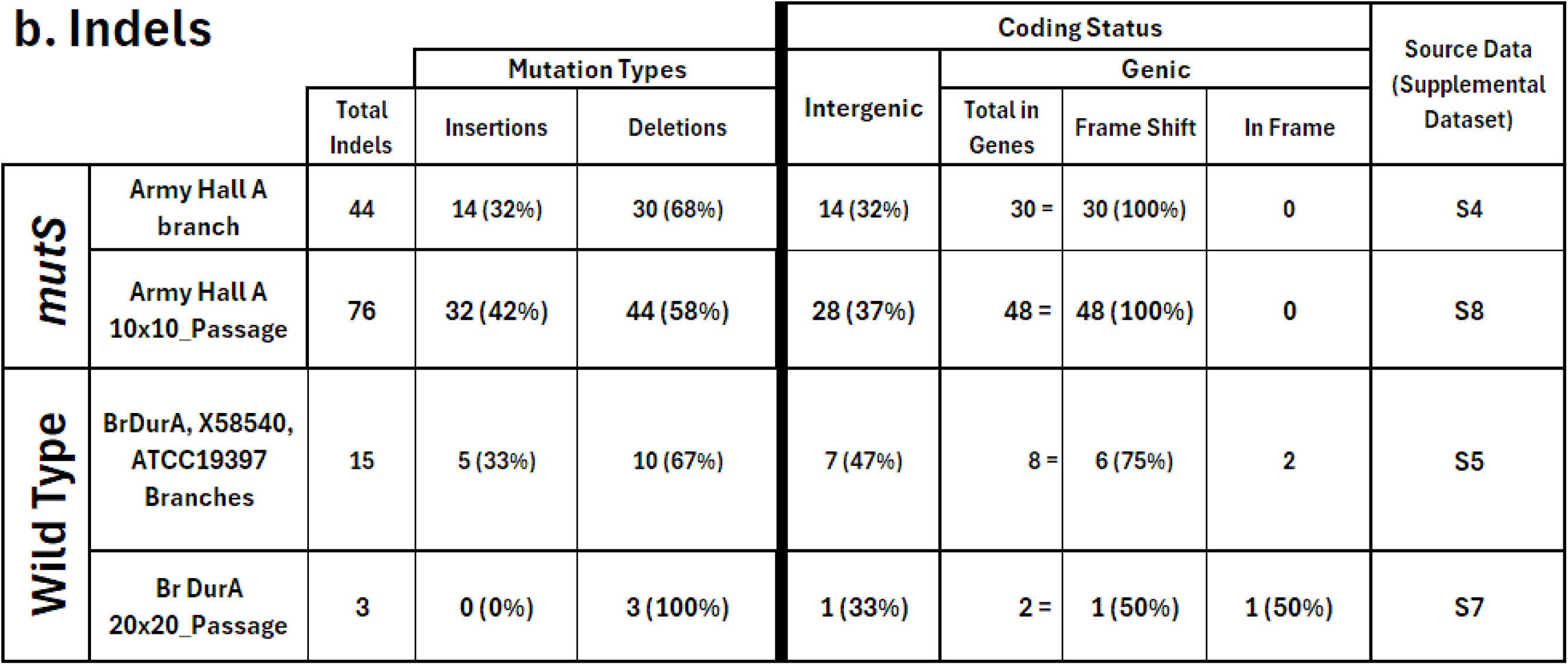
Mutational changes along phylogenetic branches and generated in experiments. **a. SNPs** found on the specified phylogenetic branches or generated during mutation-accumulation passage experiments. The branch SNPs correspond to those in Fig. 1a and the passage SNPs to those in Fig. 2. **b. Indels** found on phylogenetic branches or generated during the experiments, again corresponding to Figs. 1a and 2, respectively. The AHA strain has a *mutS* nonsense mutation while BrDuraA is a closely related wild-type strain. Precise genomic positions and annotations for each row are in the Datasets indicated.

Additionally, the AHA genome has two large multi-gene deletions, resulting in the loss of ∼82 genes (Fig. 1b). The larger deletion, involving 61 genes, may be linked to prophage excision, as 15 genes show similarity to phage genes, while the remaining 46 do not. The smaller 21-gene deletion contains only hypothetical or non-phage annotations based on amino-acid similarity. There are also two single-gene deletions in AHA (Dataset S2). Overall, the AHA genome is 55,092 nucleotides smaller than X58540, mainly due to the two large deletions (Fig. 1b, Dataset S2). Comparisons with the ATCC19397 outgroup indicate the lost genes are evolutionarily derived in AHA, with X58540 maintaining the ancestral state. This genome reduction may or may not have been adaptive under laboratory conditions (28), and the phenotypic impact is not clear from their annotation. Regardless, these genes are not essential for AHA in its new laboratory niche (29).

### Mutations identified on the phylogenetic branches

The 46 SNPs and 44 indels on the AHA branch represent the evolutionary path from nature to a laboratory-adapted bacterium (Fig. 1a). Of these 90 mutations, 82% of the SNPs and 68% of the indels are in gene-coding regions (Table 1). The genic SNPs include 27 missense, 10 synonymous, and 1 nonsense mutation (Table 1A). Several missense mutations occur in regulatory and potential signaling proteins (Dataset S3), though their effects on specific phenotypes are unknown. The single AHA nonsense mutation truncates the *mutS* open reading frame (E87X) to 9% of its full length, which likely inactivates the mismatch repair (MMR) system, thereby increasing the mutation rate and altering the spectrum of mutations. Thirty of the 44 AHA-branch indels are within genes (Table 1b). Most of these indels are single-nucleotide changes in short poly-A/T (85%) or poly-G/C (11%) tracts (*SI Appendix,* Table S1), and all of the intragenic indels cause frameshifts (Dataset S4). The affected genes include two sensory kinases and two transcriptional regulators; however, as with the missense mutations and gene deletions, their phenotypic consequences are not known. Genome decay through gene loss is common in pathogens that have altered niches and may be adaptive (28, 30, 31).

Point mutations on the AHA branch display an unusual transition-to-transversion (Ti:Tv) ratio that differs from patterns seen in wild-type strains (Table 1a) and expectations (32). Of the 46 SNPs, 41 are transitions (89%) and only 5 are transversions (11%). In comparison, the phylogenetic branches between wild-type BrDuraA, X58540, and ATCC19397 show 64% transitions and 36% transversions (Table 1a). The Ti:Tv bias on the AHA branch is consistent with a *mutS* mutation that causes a defective MMR system, one that cannot efficiently repair incipient transition mutations and thereby produces a skewed mutational spectrum (33). Another notable difference is the high frequency of indels on that same branch (Table 1b). The AHA branch has 44 indels, nearly matching the 46 SNPs. By contrast, wild-type branches have 15 indels and 42 SNPs, or about one indel for every 3 SNPs. Many indels on the wild-type branches are found in complex repeat arrays, with a smaller number in simple homomeric repeats (Dataset S5). In both cases, most indels (roughly two-thirds) occur in genic regions and all result in frameshift mutations.

### Mutations identified in replicated mutation-accumulation experiments

We performed mutation-accumulation experiments (34–36) with both AHA and BrDuraA strains in order to observe directly any differences in their rates and types of mutations (Fig. 2, Table 1). By starting each replicate lineage from a separate colony (each the outgrowth of a single cell), we ensure that observed mutations result from independent events. By serially propagating each lineage via single colonies (and thus single cells), the experimental design severely limits the power of natural selection, thereby allowing an unbiased estimation of mutation rates and the spectrum of mutational types (*SI Appendix,* Fig. S3).

**Fig. 2.**
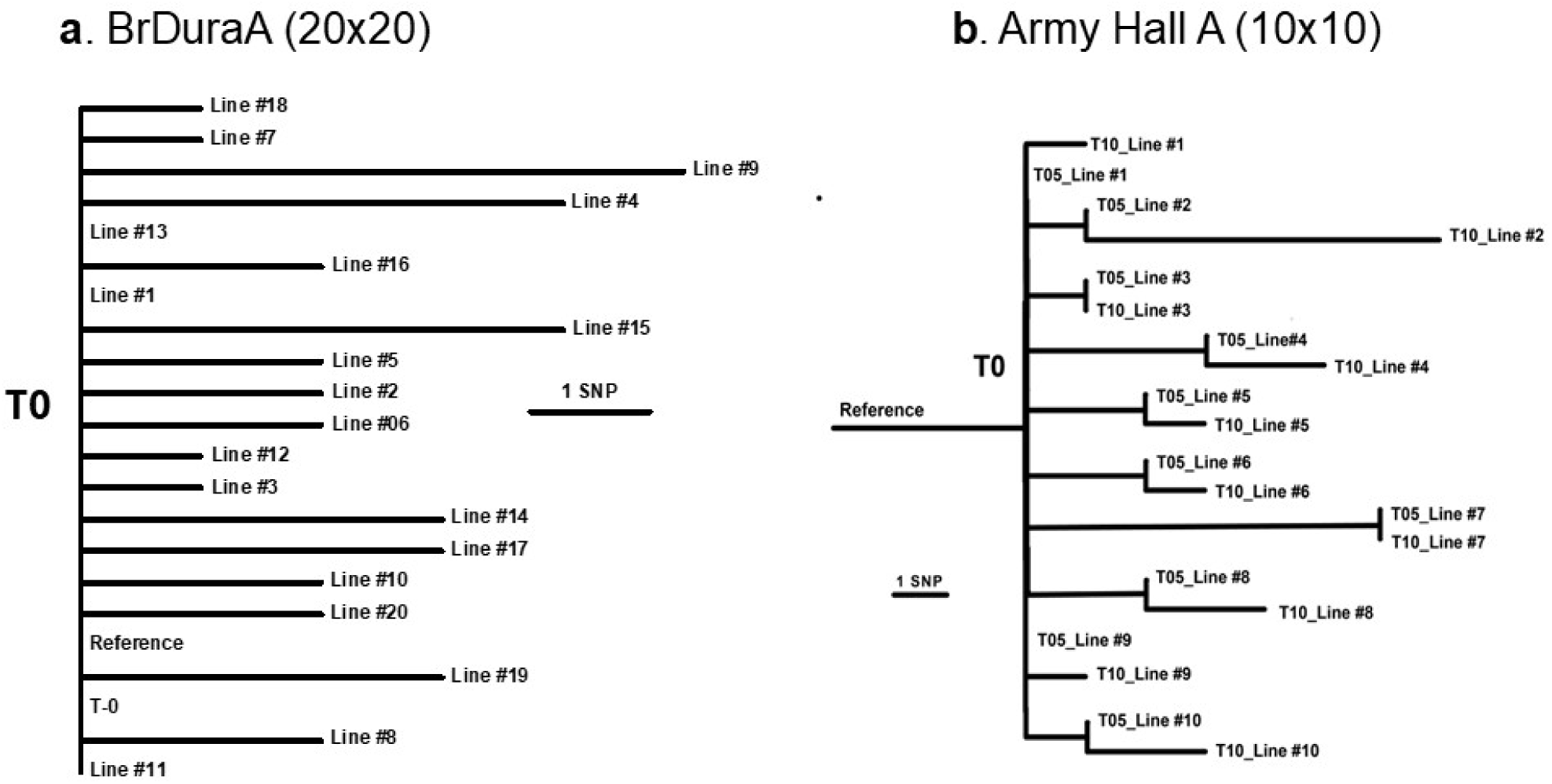
Phylogenetic summary of mutations generated in mutation-accumulation experiments. Parallel experiments were performed to generate independent lineages with known structure and generations. Whole-genome sequencing was done on endpoint (and some intermediate) samples to identify SNPs generated in each lineage. These SNPs were used to construct maximum parsimony phylogenies. **a,** Phylogeny of 20 lineages of the wild-type BrDuraA strain passaged for 20 transfers. T0 represents the BrDuraA reference genome for the initial colony used to initiate the 20 independent lineages. A total of 40 SNPs were observed after 20 transfers (Table 1a and Dataset S7), with 3 lineages having no SNPs. **b,** Phylogeny of 10 lineages of the *mutS* AHA strain passaged for 10 transfers. T0 is the AHA subclone 182-M2 (see Methods), while the CP000727.1 genome was used as the reference and differs by 7 SNPs from the T0 genome (Dataset S6). Both the T5 and T10 samples were sequenced for all 10 lineages. A total of 17 and 38 SNPs were observed at T5 and T10, respectively (Table 1a). In both experiments, the distribution of mutations appears random and fits the expectations from the Poisson distribution.

As expected from this experimental design, phylogenetic trees based on SNP data produced polytomies matching the number of replicate lines for the two strains (i.e., 10 branches for AHA and 20 for BrDuraA). In the 10 AHA lineages, a total of 38 new SNPs and 76 new indels were identified over the course of 10 passages (Table 1). The *mutS*-associated Ti:Tv bias was pronounced in AHA, with transitions accounting for 95% of the new SNPs. AHA also exhibited a high indel-to-SNP ratio, with twice as many indels as SNPs. The wild-type BrDuraA experiment was four times larger (with 20 lineages passaged for 20 transfers) and yielded 40 new SNPs (Fig. 2a), with equal numbers of transitions and transversions (Table 1). Indels were uncommon in the BrDuraA lines, with only three detected (7% of the SNPs) and only one in a homomeric repeat locus (Dataset S7).

From these data, we estimated genome-wide mutation rates given the 3.5-Mb genome size and 26.4 generations (doublings) between successive single-colony bottlenecks (see Methods). For AHA, the 38 SNPs correspond to a point mutation rate of 4.1×10^−9^ per site per generation, while the rate for BrDuraA is ∼4-fold lower at 1.1×10^−9^ per site per generation. Thus, AHA accumulates a new point mutation in roughly every 68 cell divisions, compared to every 246 divisions in BrDuraA. The distribution of intergenic, missense, synonymous, and nonsense mutations was similar for the two strains and to the patterns observed in the phylogenetic data (Table 1).

The AHA lines also had many more indels than the BrDuraA lines, again consistent with the phylogenetic data (Table 1). Both AHA and BrDuraA have ∼16,000 loci with six or more homomeric repeats (*SI Appendix,* Table S1), which are the primary sites for indels when mismatch repair is absent, as it is in the AHA strain. The 76 indels in the AHA lines occurred in both A/T (88%) and G/C (11%) tracts. Although A/T mutations were more frequent, G/C mutations occurred at a proportionally higher rate given the relative scarcity of G/C repeats (*SI Appendix,* Table S1). AHA lines also had indels in one dinucleotide repeat and one complex sequence. By contrast, the BrDuraA lines produced only 3 indels, far fewer than their 40 SNPs. The estimated indel rate is 2.9×10^−2^ per genome per generation for AHA and ∼100-fold lower at 2.8×10^−4^ for BrDuraA. Thus, AHA accumulates a new indel in roughly 35 genome replications, on average, whereas BrDuraA does so in about 3,500 replications. A single AHA colony with 10^8^ cells will contain ∼3 million indel mutations, demonstrating the dramatic effect of the *mutS* defect.

### The *mutS* mutation occurred early on the AHA branch

Determining when the *mutS* event occurred along the 46-step branch leading to AHA is important for understanding the trajectory of its domestication. However, this branch lacks internal nodes, making the order of mutations uncertain. Some useful information is nonetheless available because the *mutS* mutation alters subsequent events by strongly increasing the ratio of transition (Ti) to transversion (Tv) mutations (37, 38). The Ti:Tv ratios observed along the wild-type phylogenetic branches (64:36) and in the BrDuraA-derived experimental lines (50:50) contrast sharply with the 95:5 Ti:Tv ratio seen in the AHA lines (Table 1). Of the 46 SNPs on the AHA branch, 41 are Ti (89%) and only 5 are Tv (11%). This pronounced bias suggests the *mutS* event occurred early.

To formalize this inference, we performed a maximum likelihood analysis to estimate the probability that the *mutS* mutation was the first, second, third, etc. of the 46 events on the AHA branch. Let P and 1 – P be the relative probabilities of transversion and transition mutations in the ancestral state, and P* and 1 – P* be those probabilities after the *mutS* mutation occurred. This model corresponds to a Bernoulli process with a single change in the governing probabilities. The calculations are conditional on the observed outcome, namely that the other 45 mutations include 4 transversions and 41 transitions. The *mutS* mutation is not included in these calculations, because the loss of mismatch repair could have occurred by many different types of mutations including transversions, transitions, and indels. The analysis assumes implicitly that a near-infinitude of genomic sites could have given rise to the observed transversions and transitions.

We performed the analysis using two estimates of the wildtype transversion probability, with P = 0.5 based on the experimental data for the BrDuraA lines, and P = 0.36 using the phylogenetic data that excludes the AHA branch. We use P* = 0.05 after the *mutS* mutation based on the AHA experimental lines. In both cases, the analysis supports the inference that the *mutS* mutation was among the first 10 or so mutations on the branch leading to the AHA strain (Fig. 3). However, the true probabilities of transversions and transitions are not precisely known. Therefore, we also performed a sensitivity analysis for several combinations of P and P* that are plausible given uncertainty in their true values (*SI Appendix,* Fig. S4). These analyses confirm that the *mutS* event likely occurred early on the AHA branch, thus facilitating evolutionary change by generating genetic diversity.

**Fig. 3.**
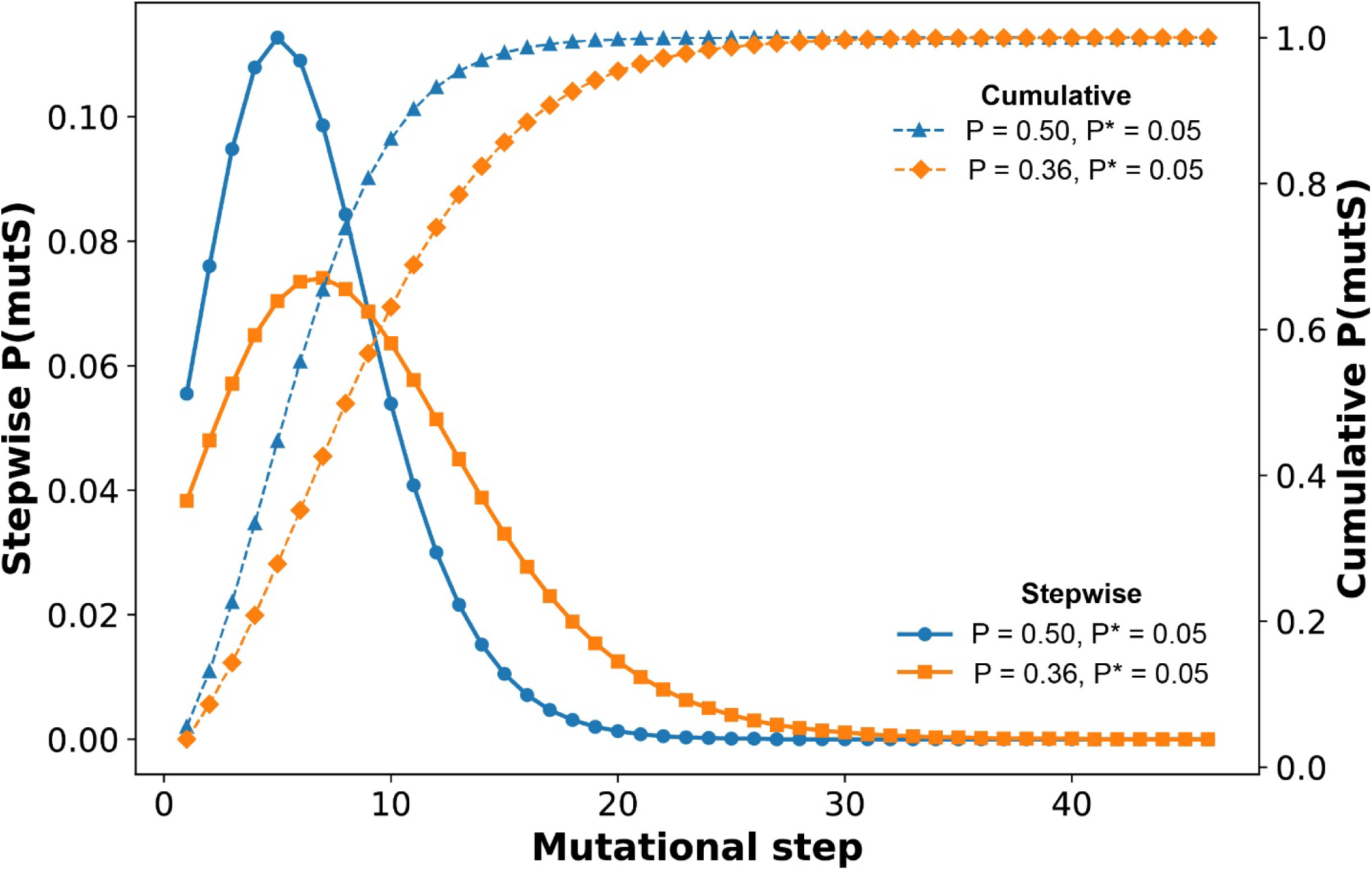
Placement of the *mutS* event on the branch leading to AHA. A maximum likelihood analysis was performed to place the *mutS* nonsense mutation on the branch leading to the AHA strain, given the change in relative frequencies of transversion and transition mutations induced by defective mismatch repair. We calculated the probability of each of the 46 possible placements of the *mutS* mutation, conditional on observing 4 transversions and 41 transitions for two ancestral probabilities of transversion (P = 0.50 and P = 0.36) using the same post-*mutS* probability of transversion (P* = 0.05) in both cases. The solid and dashed blue curves show stepwise and cumulative probabilities for P = 0.50, while the orange curves show the corresponding probabilities for P = 0.36. The stepwise probabilities for the *mutS* event peak at 5 and 7 steps, respectively, while the cumulative probabilities exceed 50% by 6 and 9 steps, respectively. The calculations for this figure are in Datasets S9a and b.

### Adaptive phenotypes in the AHA strain

The AHA and HAH strains reportedly lack the ability to form spores (17, 39). This phenotype is advantageous for industrial BoNT production and has been linked to elevated toxin yield in HAH (20). We verified the absence of spore formation in AHA by demonstrating its sensitivity to heat, with BrDuraA as a control (*SI Appendix,* Table S2). Approximately 10^8^ cells were plated for each strain, and even after only 5 min at 80°C, few AHA cells survived, and none survived 10 min of heat treatment. By contrast, BrDuraA cultures remained viable even after 30 min. Spores are critical for survival in nature, but an asporogenic variant may have a selective advantage in laboratory conditions by outcompeting wild-type strains that invest resources and delay growth by sporulating. Alternatively, the asporogenic trait may have been deliberately selected by human screening and might not provide a competitive advantage *in vitro*. We performed competition experiments to distinguish between these evolutionary scenarios.

We competed AHA against the closely related wild-type BrDuraA strain to assess AHA’s relative fitness under laboratory growth conditions. Stationary-phase cultures of each strain were mixed in three different volumetric ratios (90:10, 50:50, and 10:90) and serially transferred through ten 1/100 dilution cycles. The AHA proportions estimated by whole-genome sequencing (WGS) were slightly less at 86.8%, 42.5%, and 7.9%. Regardless, AHA quickly dominated all three mixtures, exceeding 93% abundance by the second transfer and 99% abundance by the fifth transfer (Fig. 4a). We calculated selection coefficients (s) for each mixture based on the slope of the log-transformed ratio of their relative abundance over the first transfer cycle (Fig. 4b). The AHA strain had fitness advantages of 36%, 55%, and 61% per generation for the 10:90, 50:50, and 90:10 starting ratios, respectively. However, AHA did not completely eliminate BrDuraA even after 10 transfers. Between transfers 5 and 10, the selection coefficients do not differ significantly from zero, suggesting a stable coexistence with BrDuraA persisting at low frequency. A tradeoff between growth rate and stationary-phase survival is a plausible mechanism that could sustain their coexistence. In any case, these results demonstrate that an asporogenic variant such as AHA could quickly dominate in the laboratory, even without deliberate human intervention.

**Fig. 4.**
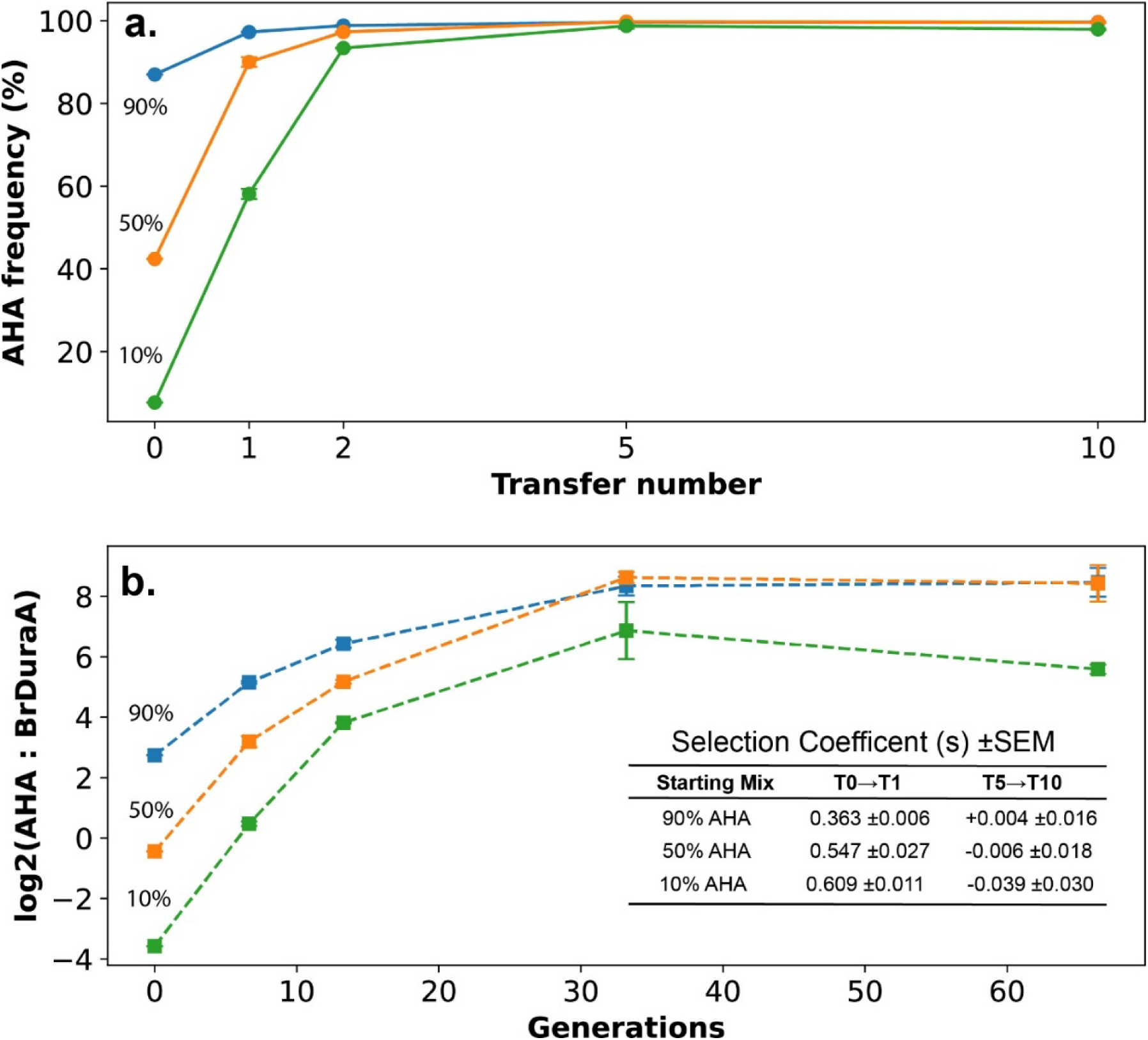
Competition experiments between the AHA and BrDuraA strains. **a,** The frequency of AHA (182-M2 subclone) was determined at five transfer points during growth in mixed culture with BrDuraA. The initial AHA:BrDuraA mixtures were 90:10 (orange), 50:50 (blue), and 10:90 (green), based upon the volumes of the respective stationary-phase cultures used to start the experiment. There were 3 replicates for each starting ratio, and the mixed cultures were transferred by 1/100 dilutions, corresponding to 6.64 cell generations, for 10 transfers. DNA was extracted at transfers 0, 1, 2, 5, and 10, and the actual relative frequencies of the two competitors were determined from the sequence data by summing read counts for distinguishing SNPs at 70 loci. **b,** The same data are shown using the log_2_ transformation of the ratio AHA:BrDuraA, which highlights the different degree of selective advantage of the AHA strain as a function of its starting frequency. The inset shows selection coefficients (s) calculated from the slopes of the curves for the three starting frequencies of AHA and over two different transfer intervals. The data for this figure are presented in Dataset S10.

## DISCUSSION

Microbial domestication is the process by which wild lineages are shaped through sustained cultivation under human influence, and it has produced many lab-adapted strains that differ in important ways from their natural ancestors (35, 40, 41). The AHA Clostridial strain has existed and proliferated for more than 90 years in laboratories, including a military toxin-production facility, resulting in a bacterium adapted to a new and restricted ecological niche. Its superior competitive fitness relative to a closely related wild-type strain under lab conditions shows the profound impact of its prolonged domestication. The asporogenic phenotype of AHA likely contributes substantially to its competitive advantage in the lab, although other metabolic and regulatory changes may also play important roles. Comparative genomic analyses reveal almost 100 genetic changes along the phylogenetic branch leading to AHA from its MRCA with wild-type lineages. None of these changes can be linked with confidence to a specific lab-adapted trait, though many are consistent with genome decay (29, 42, 43), a process often interpreted as neutral, but which in aggregate may enhance adaptation to a constant and nutrient-rich environment (44, 45). The AHA strain therefore represents an example of a wild bacterium’s domestication, offering a model for understanding the evolution of microbes under sustained cultivation by humans (40, 46, 47).

A key event in the domestication of AHA was the *mutS* nonsense mutation, which produced a hypermutator phenotype that increased the genetic diversity that fuels adaptive evolution. The domestication of many animals and plants has involved a broad gene pool and recombination, but laboratory strains typically lack these sources of diversity, amplifying the role of hypermutability. Hypermutability *per se* is not inherently adaptive and can increase the load of deleterious mutations (48, 49), but mutations that cause hypermutability can persist by hitchhiking with beneficial mutations (50). It seems improbable that the *mutS* mutation arose in nature; instead, our analysis supports its early emergence in the laboratory (Fig. 3), and its influence on subsequent genetic variation is substantial. We observed a substantial bias among SNPs toward transition mutations, along with a marked increase in indels and associated frameshift mutations leading to pseudogene formation (Table 1). Although the consequences of these loss-of-function mutations are uncertain, they may contribute to adaptive changes in cellular metabolism (51). The thousands of homomeric arrays mutate at very high rates, typically silencing genes containing them. However, this is a reversable process, in which a silenced gene can also revert at a high rate with a subsequent mutation in the same array (52). Thus, the adverse effects of Muller’s Rachet would likely be less dire for indels that can easily revert. Moreover, the evolutionary fine tuning of gene networks might be accomplished more efficiently with indels than with point mutations. The point mutation rate measured in AHA is also higher than in the wild-type comparison strain, though it is still lower than some other reported values (37). Compensatory mutations that reduce the impact of the *mutS* hypermutator phenotype have been described (53), and it is possible that a similar change occurred along the AHA branch after the *mutS* event. If so, the impact of any such compensatory mutation on indels may be small, however, as the AHA lines still generated ∼100-fold more mutations than did the wild-type lines.

In this study, we examined the domestication of the Army Hall A (AHA) strain of *C. parabotulinum*, which served as the main source of BoNT for the U.S. military and gave rise to the pharmaceutically important Hall A-hyper strain. The origins of AHA in the early 20^th^ century remain poorly documented, though Professor Ivan C. Hall is generally credited as its original source before the strain was transferred through several labs and eventually to the U.S. Army lab at Camp Detrick (Maryland) during WWII (11). This research facility was focused on biological weapons, and the WWII Allied powers regarded BoNT as both a threat to be countered and a potential offensive agent (9, 15). The Army selected AHA for its robust growth and high toxin yields during large-scale fermentation. They produced ∼1 million units to vaccinate 300,000 troops in preparation for the D-Day offensive and later for sustained defensive toxoid production (54). In the early 1970s, after the discontinuation of U.S. biological weapons programs, the AHA strain was shared with other labs, including the Food Research Institute at the University of Wisconsin. Researchers there continued to use AHA for BoNT production and coined “Hall A-hyper” (HAH) to describe its high toxin output (not its then-unknown hypermutator status), and they subsequently shared it with other research institutions and commercial entities (11). Nearly a century later, we have traced the evolutionary trajectory of this strain to the highly valuable pharmaceutical variant used extensively today.

## Materials and Methods

### *C. parabotulinum* strains

The seven species of *Clostridium* that generate neurotoxins are anaerobic, spore-forming, Gram-positive bacteria. Historically, these were all classified as *C. botulinum* but divided into Groups I through IV. As per our previous paper, we refer to the Group I strains by their species distinguishing binomials, *C. parabotulinum* and *C. sporogenes* (12). Hall strains, named for their collector Ivan Hall, are generally catalogued using a unique numeric identifier (Hall 83, Hall 11481, Hall 174, etc.); however, one particular Hall strain has come into popular usage without a numeric identifier (11). This strain has been and continues to be used by the U.S. Army, academic laboratories, and multiple pharmaceutical companies for the production of BoNT. This Hall A strain includes the “Army Hall A” (AHA) strain and its derivative, the University of Wisconsin (UW) “Hall A-hyper” (HAH) whose genome is analyzed in this paper. The HAH strain reportedly yields higher quantities of BoNT than other strains (16, 20). The UW obtained HAH from the U.S. Army program at Ft. Detrick MD in about 1970 (11), while the University of Massachusetts acquired their strain (“UMass HAH”) from UW around 1989 (55).

These strains have biosecurity risks and are controlled as Tier 1 Select Agents, which restricts their distribution and use in the U.S. only to federally licensed laboratories. An aliquot of the AHA master cell bank was generously provided by the U.S. Army Medical Research Institute for Infectious Diseases (USAMRIID) for handling at the Northern Arizona University Select Agent facility. This cell bank was the original source used to generate a single-colony subculture for the published CP000727 genome (56). From this AHA master cell bank, we established a single-colony stock (182-M2) for the passage experiment (Fig. 2b) and to verify the sequences in the CP000727.1 FASTA accession (Dataset S3). The CP000727.1 genome has become a research standard, but it was generated by Sanger sequencing and released without raw data, making verification of its accuracy problematic. Using the 182-M2 genome we found only 7 SNPs different from the CP000727 sequence, which we document in Dataset S6 and show as the basal branch in Fig 3b. The Wisconsin HAH strain DNA was provided by Dr. Sabine Pellett at UW and sequenced at NAU.

The BrDuraA strain is phylogenetically the closest strain to AHA in the NAU inventory; it was kindly provided by Prof. Rafael A. Fernández (Universidad Nacional de Cuyo, Mendoza, Argentina). BrDuraA was isolated in the 1980s from a case of foodborne botulism associated with canned peaches in syrup. “Br” is an abbreviation for “brote” (meaning outbreak in Spanish) and “Dura” is short for “duraznos en almibar” (peaches in syrup). BrDuraA has an A1 toxin serotype and is closely affiliated with the AHA clade (*SI Appendix,* Fig. S1b). Its genome differs from the MRCA to AHA and X58540 by only 31 SNPs (Fig. 1a) and represents the best available control strain for our studies. The BrDuraA genome was completed using a combination of short-read Illumina and long-read Oxford Nanopore data. The X58540 strain was isolated in 2005 from cattle feces (27), and its high-quality genome was recently deposited at NCBI by a South Korean company (Chong Kun Dang Bio Co Ltd.). The live strain no longer exists in Germany and is unavailable from Korea due to proprietary and biosecurity restrictions (personal communication, Frank Gessler).

### Whole-genome sequences

All genomes used in this study are available at NCBI (*SI Appendix,* Fig. S1 and Table S1) or for the newly generated genomes AHA 182-M2 (SAMN50994695) and UW-HAH (SAMN51230428). The 182-M2 genome data was from a single-colony isolate from the Army’s master cell bank and used to verify genotypic calls of the CP000727 reference genome, which was generated by Sanger sequencing methods over 20 years ago.

### Calculating MASH distances and phylogenetic analysis

Genomic data representing *C. botulinum* and *C. sporogenes* were downloaded from NCBI databases. All available genome assemblies were downloaded from GenBank (August 2025) as well as read data for 136 entries from the SRA database (105 Illumina paired-end reads, 31 Ion Torrent single-end reads). Illumina reads were trimmed with Trim Galore v0.6.10 (--cores 4 –paired) (https://github.com/FelixKrueger/TrimGalore), which uses Cutadapt v4.9 (57) and FastQC v0.12.1 (https://www.bioinformatics.babraham.ac.uk/projects/fastqc/), and assembled with the SPAdes genome assembler v4.0.0 (--isolate -t 4 -m 16 --cov-cutoff auto) (58). Ion Torrent reads were assembled with the SPAdes genome assembler v4.0.0 (--isolate --iontorrent -t 4 -m 16 --cov-cutoff auto). MASH (version mash-Linux64-v2.3) (59) distances between all genomes were computed using a sketch size of 1000 and a kmer size of 21. Any genome with a MASH distance greater than 0.08 (∼92% average nucleotide identity) from the Army Hall A genome (GCA_000017045.1/CP000727.1) was excluded from the dataset to focus the analysis on *C. parabotulinum* and *C. sporogenes* (12). A neighbor-joining tree for the remaining 1104 genomes was generated from the MASH distance matrix with phylip (v3.697) (60, 61). The tree was midpoint rooted, which separates the two Group I species, *C. parabotulinum* and *C. sporogenes*.

### Genome alignment and visualization

Genomic comparisons and visualizations were generated with pyGenomeViz v1.2.1 (https://github.com/moshi4/pyGenomeViz) using either the progressiveMauve (snapshot_2015_02_13) (62) (Fig. 1b) or MUMmer v3.23 (63) (*SI Appendix,* Fig. S2) alignment options.

### Single nucleotide variant discovery and phylogenetic analysis

Paired-end Illumina reads were aligned against relevant references with minimap2 v2.29 (64) and single nucleotide polymorphisms (SNPs) were identified with GATK v4.5.0 (65, 66). For genome assemblies where raw data were not available, reads were simulated from the assembly with ART vART-MountRainier-2016-06-05 (67). SNPs that fell within duplicated regions of the genome, based on a reference self-alignment with NUCmer v3.23 (68), were filtered from downstream analyses. SNP positions were also filtered if the depth of read coverage was <5 or the allele call proportion was <0.9. All of these methods were wrapped by NASP v1.2.1 (69). For outgroup rooting, SNPs were queried from an outgroup with a custom script (https://gist.github.com/jasonsahl/d1472d34198e857ed6594d3505cf55b9). The annotation of SNPs was performed with snpEff v5.1f (70). Indels were identified with snippy v4.6.0 (https://github.com/tseemann/snippy).

Maximum likelihood phylogenies were inferred on concatenated SNP alignments with IQ-TREE v2.3.6 (71) along with the integrated ModelFinder method (72). Maximum parsimony phylogenies were inferred on concatenated SNP alignments with Phangorn v2.11.1 (73).

### Two approaches to characterizing wildtype and *mutS* mutations

We examined the effect of the AHA *mutS* defect on the rate and spectrum of mutations by identifying genomic changes on phylogenetic branches and by performing a mutation-accumulation experiment to observe the generation of new mutations under controlled conditions. Retrospectively, we examined genetic variation on the phylogenetic branch between AHA (with its *mutS* defect) and its MRCA with X58540, and we then compared these data with the wild-type branches among ATCC19397, X58540 and BrDuraA (Fig. 1A). Prospectively, we propagated replicate lines of the AHA and BrDuraA strains (Fig. 2) through single-cell bottlenecks to observe the accumulation of new mutations (*SI Appendix,* Fig. S3).

The mutation-accumulation experiments used the *mutS* AHA subclone 182-M2 and the closely related *mutS*-wildtype strain BrDuraA. A single colony of each was used to start 10 independent *mutS* lineages and 20 independent BrDuraA lineages. A single colony was chosen at random to propagate each lineage at each transfer, with 10 transfers for the AHA lines and 20 for the BrDuraA lines. Each colony was suspended in 0.5 mL of broth and then used for the next transfer, frozen with 15% glycerol for storage, and used for DNA isolation at transfers T0, T5, and T10 for AHA, and T0 and T20 for BrDuraA.

Three other typical colonies were harvested and titered to estimate the number of CFUs in each. The average CFU estimate was 87.4 million (s.d. = 27.6 million). On average, each colony thus required log_2_ (8.74 x 10^7^) ≈ 26.4 doublings to reach its size at each transfer. Given a queryable genome of 3.5 million nucleotides, the 10 x 10 experiment with AHA involved ∼9.2 billion nucleotide replication events, and the 20 x 20 experiment with BrDuraA ∼37 billion events. These numbers were used to estimate corresponding mutation rates.

### Inferring the placement of the *mutS* event on the AHA branch

Defects in mismatch repair, such as those caused by the *mutS* nonsense mutation on the phylogenetic branch leading to the AHA strain, increase the frequency of transition mutations relative to transversion mutations. There are 46 point mutations on the AHA branch. To infer the likelihood of each placement of the *mutS* mutation among the 46 events on the AHA branch, we model a Bernoulli process with a single change point. Let P and 1 – P be the probabilities of transversion and transition mutations in the ancestral state, and P* and 1 – P* be those probabilities after the *mutS* mutation. The likelihood calculations are conditional on the observed outcome, namely that the 45 other mutations include 4 transversions and 41 transitions. (The *mutS* mutation is not included because the loss of mismatch repair could have occurred by many different types of mutations including transversions, transitions, and indels.) We also assume sampling with replacement because a near-infinitude of genomic sites could have given rise to the observed transversions and transitions.

The likelihood for the mutator at position m is given by the following three equations with these terms: *m* = position of the mutator on the phylogenetic branch, *n* = number of mutations that occur before the mutator (*m*-1), v = number of transversions (4 in our example), *s* = number of transitions (41 in our example), *p* = probability of a transversion before mutator, *p\** = probability of a transversion after mutator, (*1-p*) = probability of a transition before mutator, (*1-p\**) = probability of a transition after mutator.

The likelihood for the mutator at position m is given by the following equations:

(1) For: s ≥ *n* ≥ *v*

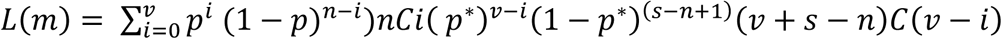
(2) For: *v* > *n* ≤ *s*

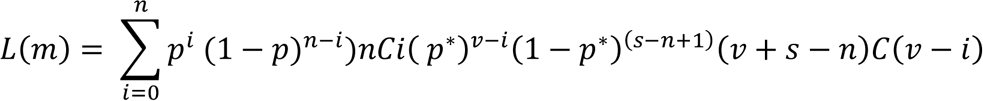
(3) For *v* < *n* > *s*

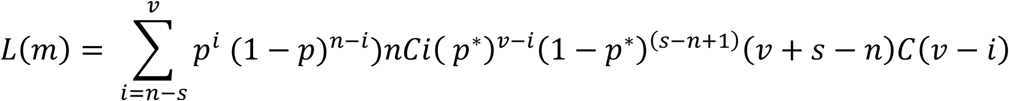
(4) The probability of the mutator at position *m* is then given by:

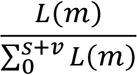

Fig. 3 shows the stepwise and cumulative probabilities for the placement of the *mutS* event using estimates for P and P* from the experimental mutation-accumulation lines, as well as for P from other phylogenetic branches. *SI Appendix,* Fig. S4 presents a sensitivity analysis, in which P and P* are increased or decreased moderately relative to those point estimates, and which supports the inference that the *mutS* event occurred early on the AHA branch. We also provide a spreadsheet in Dataset S9 that allows the user to enter values for P and P*, which then calculates the corresponding likelihoods for every possible placement of the *mutS* event on the phylogenetic branch leading to AHA. This spreadsheet was used to test the model’s sensitivity to differences in parameters (*SI Appendix,* Fig. S4).

### Competition experiment

We performed competition experiments to estimate the fitness of the AHA strain relative to its close relative, BrDuraA. Each strain was grown from a frozen stock in 3 mL TPGY broth. The two competitors were mixed at three different volumetric ratios (90:10, 50:50, and 10:90), which were then split into three replicates (9 cultures total). These competition cultures were transferred 10 times with 1/100 dilutions (0.03 ml into 2.97 ml of fresh broth). The starting inoculations were ∼10^7^ CFU, thus avoiding bottleneck effects. Each culture was grown anaerobically for 48 h at 35°C between transfers. We sequenced the AHA and BrDuraA stocks, the three T0 mixtures, and the 9 cultures at T1, T2, T5, and T10 (41 total). The ratio of the competitors in each sample was estimated by summing the read counts at 70 SNP loci that differed between the two strains (Dataset S10), with >50,000 reads used to estimate the AHA:BrDuraA ratio in each sample. In all, over 2 million sequencing reads were aligned across the 70 SNP positions and base calls differing from the either AHA or BrDuraA alleles were used to estimate an error rate of 0.18% per read per position. This rate is consistent with our previous studies and while low, it would have an outsized effect when estimating the frequency of minor alleles as the dominant allele “bleeds over.” This issue became important at later timepoints in the competition experiment and, hence, was used to correct the data for Fig. 4.

## Data availability

AHA assembly and passage data have been deposited at NCBI under biosamples SAMN50994695 (AHA), SAMN51230428 (HAH), and SAMN07185497 (BrDuraA). The annotated genome sequences of the AHA and BrDuraA are available at NCBI (JBROGD00000000). Gene deletions, SNPs, and indels are extensively documented in the supplementary Datasets.

## Acknowledgments

The authors would like to thank the United States Army Institute for Infectious Disease for providing the Army Hall A strain for this study. All experiments using live *C. botulinum* strains were conducted in the select agent facilities at Northern Arizona University and the University of Wisconsin-Madison. We also thank Ms. Amber Jones for the genome sequencing, which was performed at the Pathogen and Microbiome Institute. We thank Ms. Jenny M. B. Keim for assistance in the *mutS* event modeling studies. The work was funded by the E. Raymond and Ruth Reed Cowden Endowment for Food Microbiology at Northern Arizona University. R.E.L. acknowledges support from AgBioResearch (Hatch project MICL02839) and the John A. Hannah endowment at Michigan State University. Work conducted in the Pellett lab at UW-Madison was funded by the Division of Intramural Research, National Institute of Allergy and Infectious Diseases (R01 AI139306).

## Author Contributions

P.K., R.G.P., S.P., and J.W.S. supervised the project. P.K., J.W.S., and R.E.L. conceptualized the project. P.K., R.N., A.J.V., S.P., and R.E.L. designed experiments. R.N., M.A.G., and A.J.V. conducted the mutation-accumulation experiments and DNA extractions. C.H.D.W., P.K., and J.W.S. analyzed the DNA sequences, performed phylogenetic analyses, and genome alignments. E.F.M., R.G.P., R.E.L., and P.K. performed mathematical modelling of the *mutS* probability. P.K., R.E.L., and T.S. wrote the original draft of the paper. P.K., R.E.L., J.W.S., R.N., A.J.V., S.P., M.A.G., C.H.D.W., R.E.P., T.S., and E.F.M. edited the paper. P.K. and S.P. obtained funding for the project.

***Supplemental Information Appendix*** (single pdf file)

- *SI Appendix,* Fig. S1. Phylogenetic structure of the *C. botulinum* Group I strains based on whole-genome sequences.
- *SI Appendix,* Fig. S2. The AHA-specific translocation.
- *SI Appendix,* Fig. S3. Design of mutation-accumulation experiments.
- *SI Appendix,* Fig. S4. Parameter sensitivity of the *mutS* probability analysis.
- *SI Appendix,* Table S1. Homomeric nucleotide array frequency and indels associated with the *mutS* mutation.
- *SI Appendix,* Table S2. The Army Hall A strain is asporogenic.

**Supplemental Datasets (**multi-tab Excel Spreadsheet)

- Dataset S1a. *Clostridium botulinum* Group I genomes for the phylogeny shown in *SI Appendix,* Fig. S1a
- Dataset S1b. SNPs used to construct the phylogeny shown in *SI Appendix,* Fig. 1b
- Dataset S2a. Chromosomal deletions on or relative to the Army Hall A branch
- Dataset S2b. Gene annotations for the Army Hall A genome and their occurrence in related strains
- Dataset S2c. Annotation for Army Hall A gene deletions
- Dataset S3. Army Hall A branch SNPs
- Dataset S4. Army Hall A branch indels
- Dataset S5. SNPs and indels on the wildtype BrDuraA, X58540, and ATCC19397 branches
- Dataset S6. SNPs from the Army Hall A 10 x 10 passage experiment
- Dataset S7. SNPs and indels from the BrDuraA 20 x 20 passage experiment
- Dataset S8. Indels from the Army Hall A 10 x 10 passage experiment
- Dataset 9a. Maximum likelihood modeling of *mutS* event placement with P = 0.50 and P* = 0.05
- Dataset S9b. Maximum Likelihood modeling of *mutS* event placement with P = 0.50 and P* = 0.05
- Dataset S10. Read count data from 70 SNP loci for estimating AHA percentage in mixed competition cultures

## SI Appendix

**Fig. S1.**
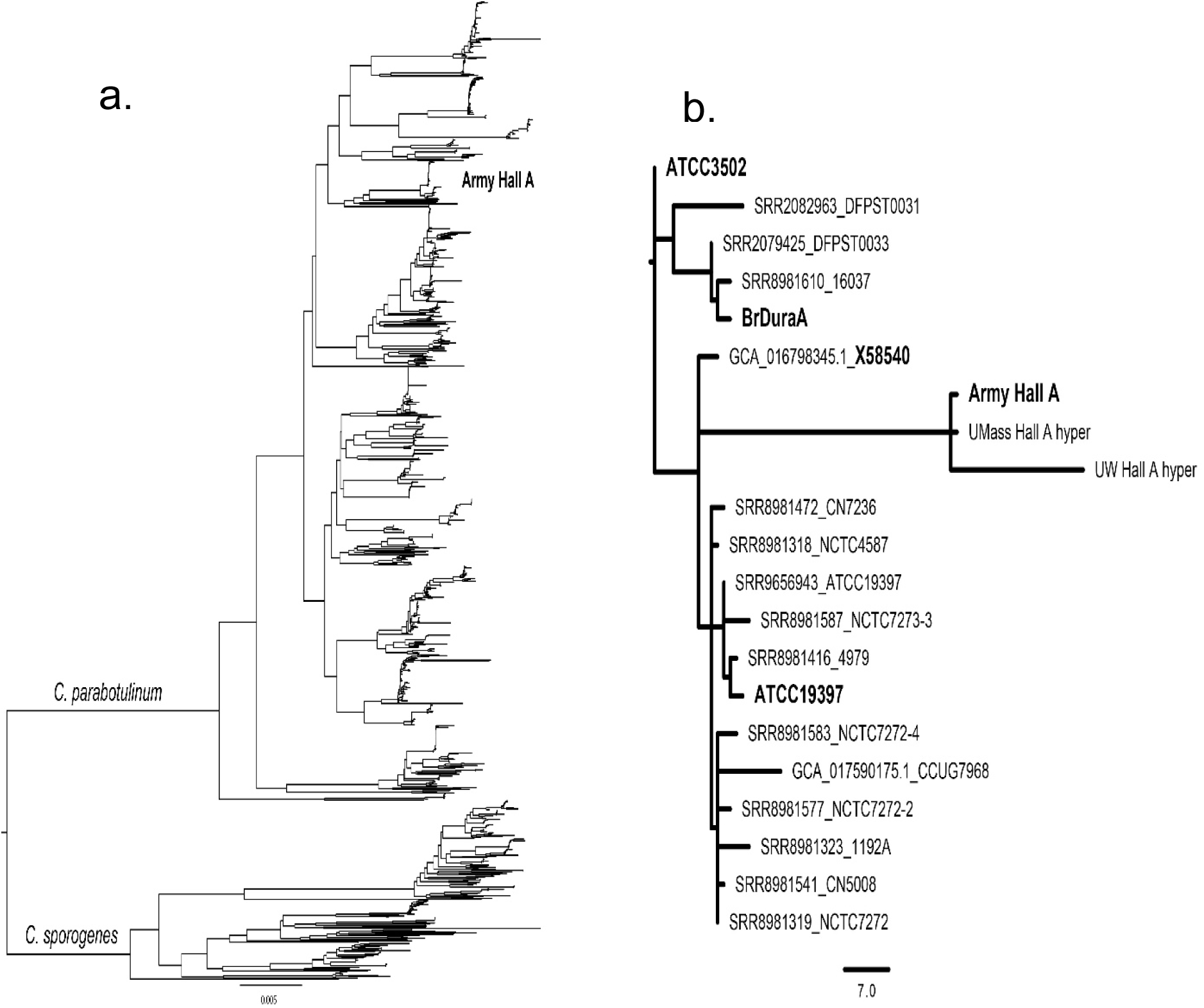
The phylogenetic structure of *C. botulinum* Group I strains based on whole-genome sequences. **(a)** Neighbor-joining tree based on minhash distances among 1104 genomes (Datset S1a). The Group I strains fall into the *C. parabotulinum* and *C. sporogenes* species, consistent with previous reports^1,2^. The location of the Army Hall A (AHA) strain within *C. parabotulinum* is indicated and expanded in panel b. **(b)** Maximum-parsimony analysis of 20 genomes from strains closely related to AHA based on 135 SNPs (Dataset S1b), rooted with the ATCC3502 genome. This group of mostly laboratory strains appears clonal, with no homoplasies (consistency index = 1). The AHA-derived HAH genomes from the University of Massachusetts (UMass) and the University of Wisconsin (UW) are on the same long branch as AHA.

**Fig. S2.**
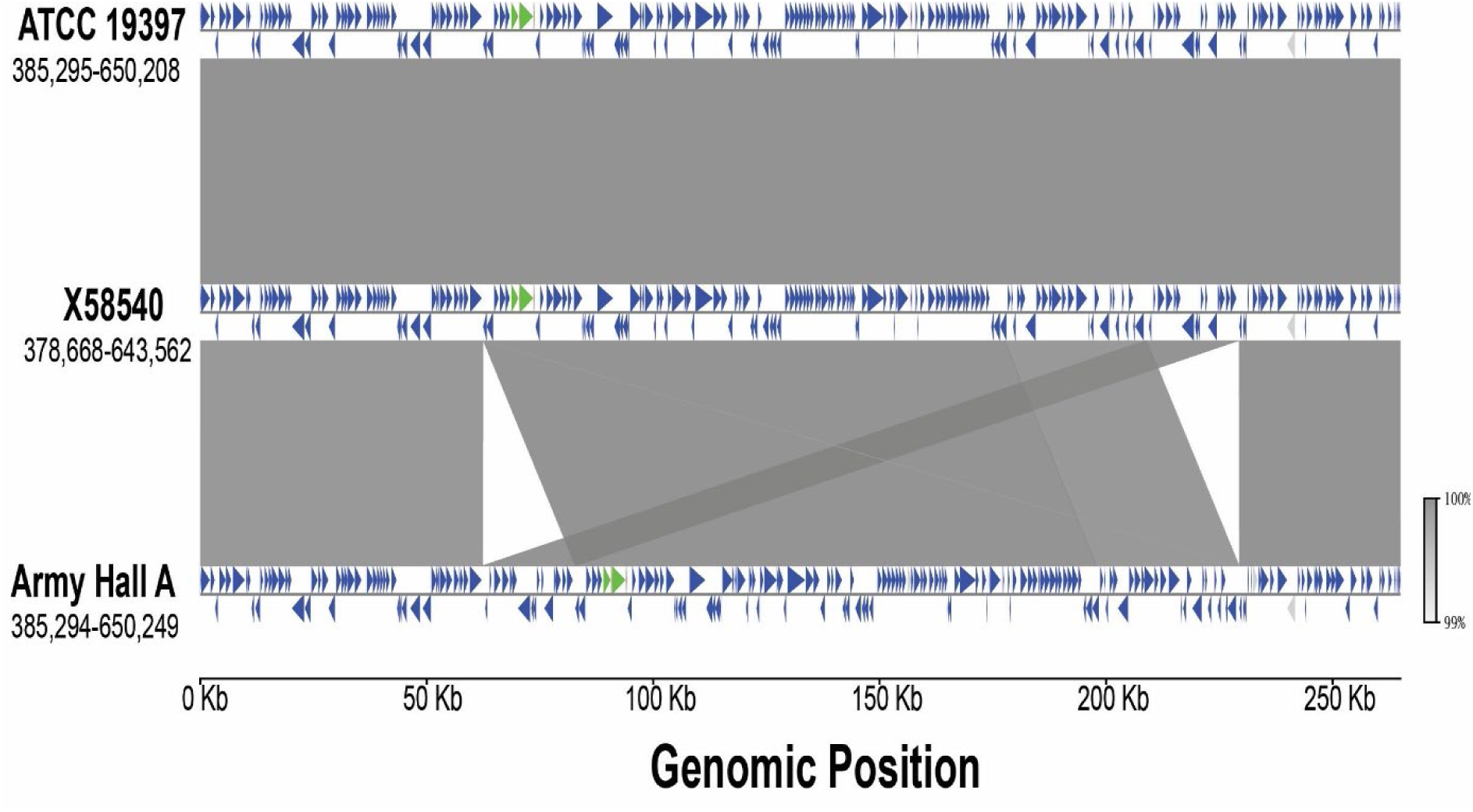
The AHA-specific translocation. The genomic regions associated with the AHA-specific translocation^3^ were aligned using MUMmer^4^. Open reading frames are shown by blue triangles that indicate their strand orientation. Green triangles represent ribosomal RNA genes. The ATCC19397 genome serves as an outgroup, which demonstrates that the translocation is ancestral in X58540 and derived in Army Hall A (CP000727). The genomic coordinates for the three aligned segments are shown below each strain label. An HTML version (https://github.com/chawillia/Cbotulinum_domestication) of this figure is available that allows interactive exploration of the genes and their annotations.

**Fig. S3.**
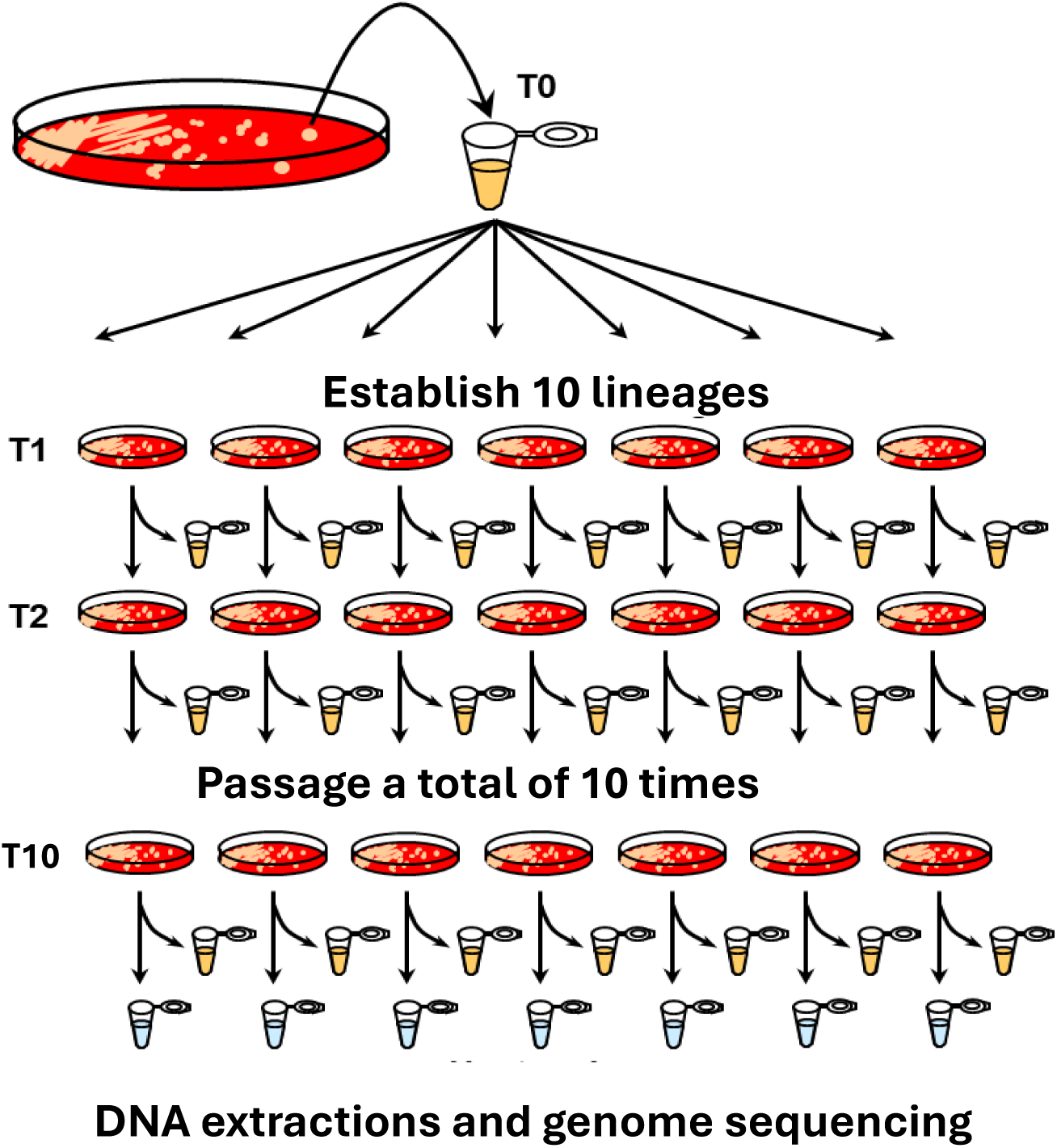
Design of mutation-accumulation experiments. This diagram illustrates the experimental workflow for the results shown in Fig. 3. A single colony was taken from an isolation streak of the AHA strain, and similarly for the BrDuraA strain. That colony was dispersed into broth (T0) and then re-streaked onto plates to generate single colonies. For AHA,10 colonies were picked to establish 10 parallel lineages (lines 1-10), which were transferred via single colonies 10 times (T1-10), with cells retained for whole-genome sequencing at each step. The process was identical for BrDuraA except that 20 lines were established and transferred 20 times each. Each colony is derived from a single cell; passaging each lineage via a single colony leads to a genetic bottleneck that fixes or eliminates any variants that occurred during that colony’s growth. About 26.4 doubling cell generations occur between transfers (see Methods).

**Fig. S4.**
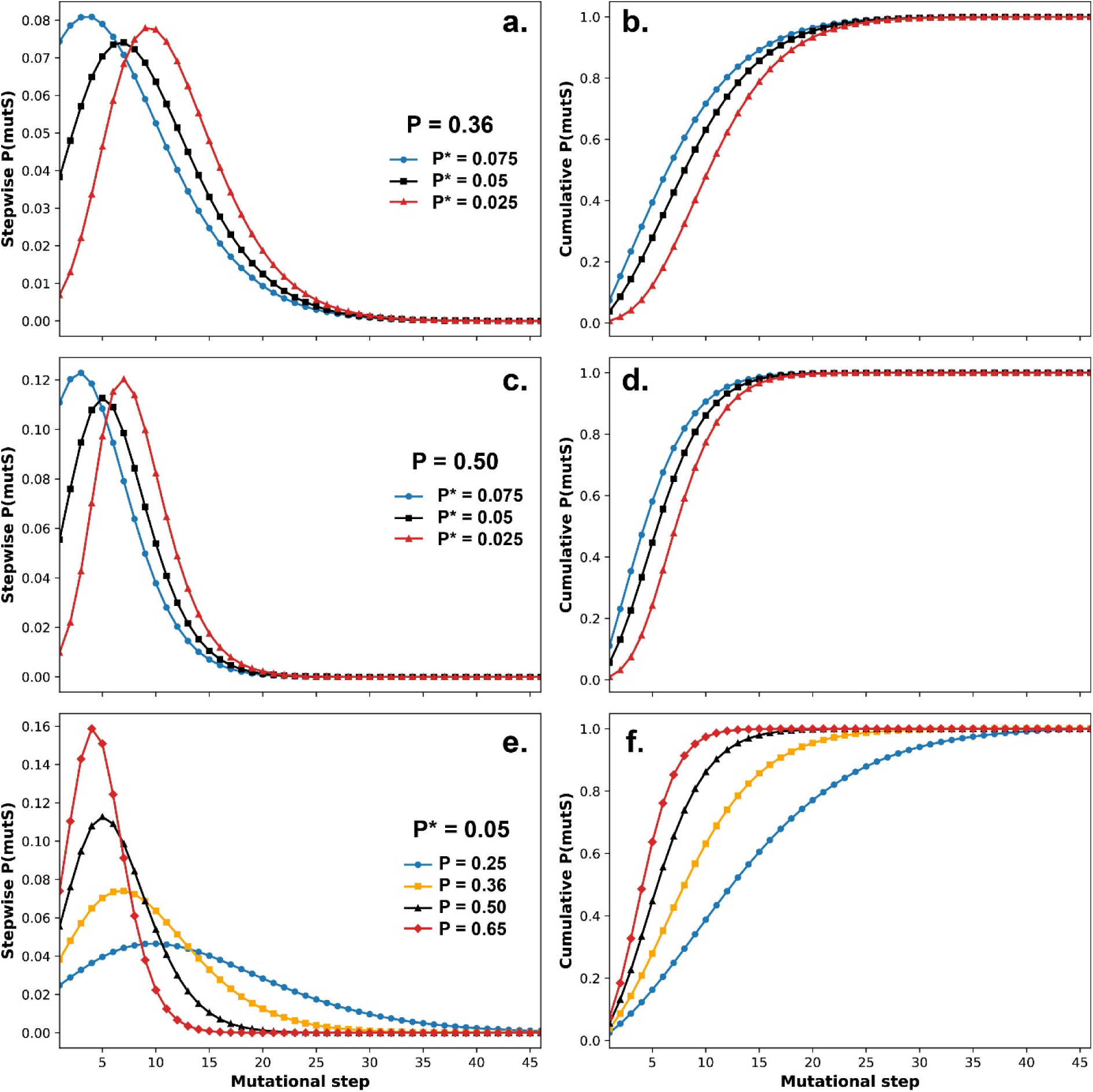
Parameter sensitivity of the *mutS* probability analysis. We examined the sensitivity of the placement of the *mutS* nonsense mutation on the 46-step branch leading to the AHA strain by changing the transversion probabilities before (P) and after the *mutS* event (P*) in the maximum likelihood analysis. Stepwise and cumulative P(*mutS*) distributions were calculated for the observed values of P = 0.36 (**a**, **b**) and P = 0.5 (**c**, **d**) with three different P* values (0.075, 0.05, and 0.025). (**e**, **f**) The P(*mutS*) distributions were also calculated for the observed value of P* = 0.05 with four different P values (0.25, 0.36, 0.5, and 0.65). The distributions were heavily skewed to early steps in all cases, supporting the conclusion that the *mutS* nonsense mutation occurred early on the branch leading to AHA.

**Table S1.** Homomeric nucleotide array frequency and indels associated with the *mutS* mutation. The numbers of homomeric nucleotide arrays in wild-type BrDuraA and *mutS* AHA (CP000727) genomes were enumerated for A/T and G/C repeats. The reported indels were observed on the AHA phylogenetic branch or in the AHA mutation-accumulation experiment (Dataset S4 and S8). There is no significant difference in the number or size of the homomeric repeats between the two strains; the minor differences reflect small differences in their core genomes, not extensive mutations between these close relatives (see Table 1). In these A+T rich genomes, the A/T arrays were larger and more numerous than the G/C arrays. The longer arrays had a higher frequency of indels, though the larger number of smaller arrays contained most of the observed mutations observed (e.g., 37 indels for 7-mers). As homomeric array size increases beyond 8, there is a slight decline in the fraction found in gene encoding regions. The proportion of homomeric repeats in genes is higher than the proportion of indels observed in genes (Table 1), which suggests that there is selection against frameshift mutations in most genes.

|  | BrDuraA |  |  | Army Hall A |  |  |  |  |  |  |
| --- | --- | --- | --- | --- | --- | --- | --- | --- | --- | --- |
| Array Size (bp) | A/T | G/C | Total Repeat Loci | A/T | G/C | Total Repeat Loci | % in coding regions | Indels* | % of indels | Indel Freq/locus |
| 15 | 0 | 0 | 0 | 0 | 0 | 0 | -- | 0 | 0% | -- |
| 14 | 0 | 0 | 0 | 0 | 0 | 0 | -- | 0 | 0% | -- |
| 13 | 1 | 0 | 1 | 1 | 0 | 1 | 0% | 1 | 1% | 1.0000 |
| 12 | 1 | 0 | 1 | 0 | 0 | 0 | -- | 0 | -- | -- |
| 11 | 1 | 0 | 1 | 1 | 0 | 1 | 0% | 1 | 1% | 1.0000 |
| 10 | 6 | 0 | 6 | 8 | 0 | 8 | 63% | 6 | 4% | 1.0000 |
| 9 | 68 | 0 | 68 | 64 | 2 | 66 | 61% | 12 | 11% | 0.1765 |
| 8 | 970 | 5 | 975 | 940 | 4 | 944 | 79% | 28 | 26% | 0.0287 |
| 7 | 4,234 | 21 | 4,255 | 4,180 | 19 | 4,199 | 75% | 37 | 31% | 0.0087 |
| 6 | 11,138 | 129 | 11,267 | 11,004 | 129 | 11,133 | 75% | 24 | 21% | 0.0021 |
| 5 | 26,630 | 1,019 | 27,649 | 26,331 | 998 | 27,329 | 75% | 6 | 5% | 0.0002 |
| total = | 43,049 | 1,174 | 44,223 | 42,529 | 1,152 | 43,681 |  | 115 |  |  |

**Table S2.**
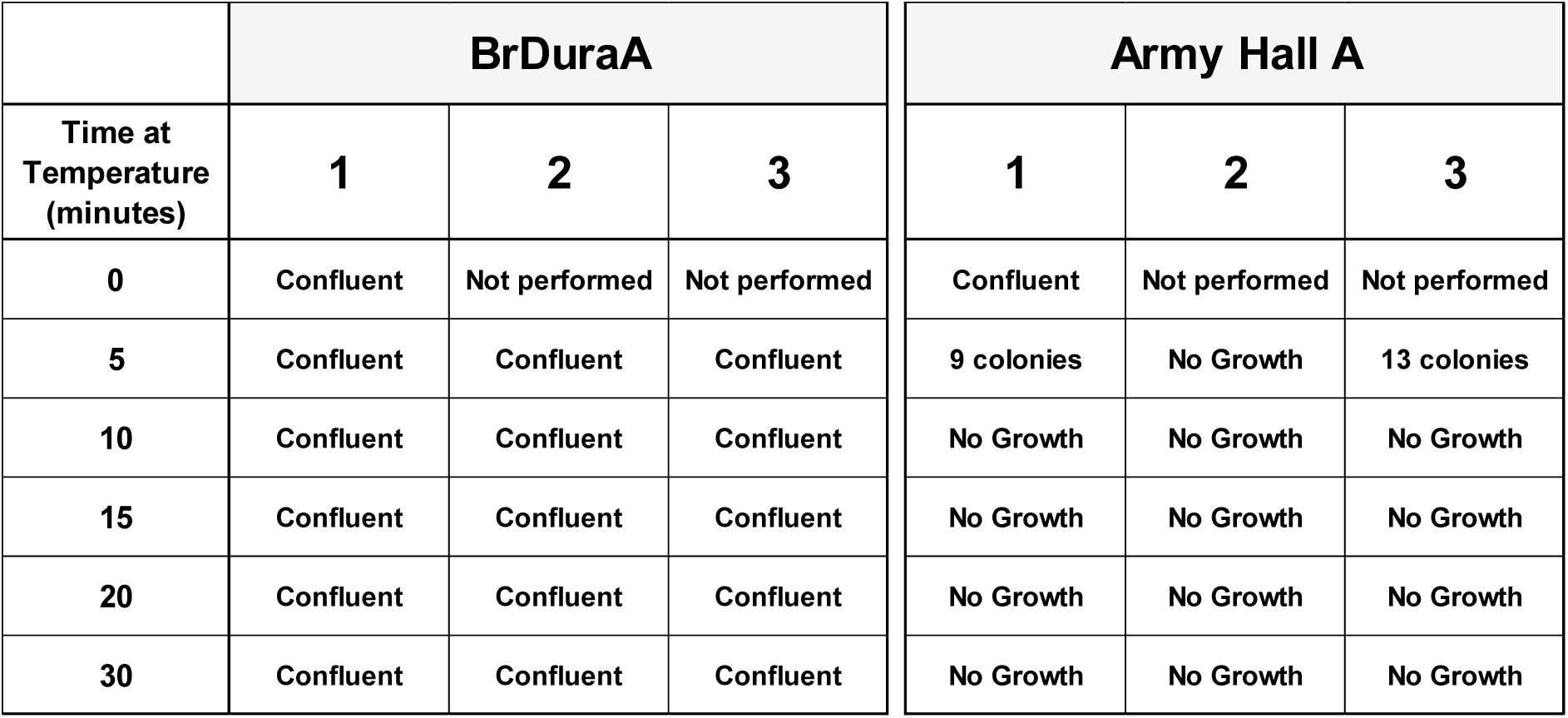
The Army Hall A strain is asporogenic. The BrDuraA strain and the Army Hall A strain were grown to stationary phase and then treated with heat at 80°C for up to 30 min. Aliquots (200 ul, ∼10^8^ CFU) were plated and incubated for 3 days to determine viability, which was scored visually. Prior to heat treatment, both strains produced confluent growth. The wild-type BrDuraA strain showed confluent growth even after 30 min of heat treatment, consistent with robust sporulation, survival, and growth. By contrast, the AHA strain produced few survivors even after just 5 min of heat treatment, and none after 10 min, indicating that it does not produce spores under these standardized conditions. The AHA cells in this table were directly from the master cell bank obtained from USAMRIID and represent the ancestral repository for the CP000727 and 182-M2 subclones used in this paper, as well as the UW Hall A-hyper cells. These three derived subclones have been shown to be asporogenic, as well. There are no spores produced in these cell banks under these standard growth conditions – not even at a low level as a minor component.

|  | BrDuraA |  |  | Army Hall A |  |  |
| --- | --- | --- | --- | --- | --- | --- |
| Time at Temperature (minutes) | 1 | 2 | 3 | 1 | 2 | 3 |
| 0 | Confluent | Not performed | Not performed | Confluent | Not performed | Not performed |
| 5 | Confluent | Confluent | Confluent | 9 colonies | No Growth | 13 colonies |
| 10 | Confluent | Confluent | Confluent | No Growth | No Growth | No Growth |
| 15 | Confluent | Confluent | Confluent | No Growth | No Growth | No Growth |
| 20 | Confluent | Confluent | Confluent | No Growth | No Growth | No Growth |
| 30 | Confluent | Confluent | Confluent | No Growth | No Growth | No Growth |

## References

1. G. Larson et al., Current perspectives and the future of domestication studies. Proc Natl Acad Sci U S A 111, 6139–6146 (2014).

2. A. T. Lin, R. A. Fairbanks, J. Barba-Montoya, H. L. Liu, L. Kistler, A legacy of genetic entanglement with wolves shapes modern dogs. Proc Natl Acad Sci U S A 122, e2421768122 (2025).

3. A. H. Freedman et al., Genome sequencing highlights the dynamic early history of dogs. PLoS Genet 10, e1004016 (2014).

4. M. D. Purugganan, D. Q. Fuller, The nature of selection during plant domestication. Nature 457, 843–848 (2009).

5. B. Gallone et al., Domestication and Divergence of Saccharomyces cerevisiae Beer Yeasts. Cell 166, 1397–1410 e1316 (2016).

6. D. Charlesworth, J. H. Willis, The genetics of inbreeding depression. Nat Rev Genet 10, 783–796 (2009).

7. J. Steensels, B. Gallone, K. Voordeckers, K. J. Verstrepen, Domestication of Industrial Microbes. Curr Biol 29, R381–R393 (2019).

8. M. Pirazzini, O. Rossetto, R. Eleopra, C. Montecucco, Botulinum Neurotoxins: Biology, Pharmacology, and Toxicology. Pharmacol Rev 69, 200–235 (2017).

9. S. S. Arnon et al., Botulinum toxin as a biological weapon: medical and public health management. JAMA 285, 1059–1070 (2001).

10. I. J. Pflug, Science, practice, and human errors in controlling Clostridium botulinum in heat-preserved food in hermetic containers. J Food Prot 73, 993–1002 (2010).

11. T. J. Smith, The Many Journeys of Botulinum Neurotoxins and the Bacteria that Produce Them. Microbiology and Molecular Biology Review 89, 00–00.

12. T. Smith, C. Williamson, K. Hill, J. Sahl, P. Keim, Botulinum neurotoxin-producing bacteria. Isn’t it time that we called a species a species? mBio 9:e01469–18., supplementary information (2018).

13. E. S. McCoy (1943) Unpublished.

14. C. Lamanna, E. O. Mc, H. W. Eklund, The purification and crystallization of Clostridium botulinum type A toxin. Science 103, 613 (1946).

15. J. L. Middlebrook, and D.R. Franz, "Botulinum Toxins" in Medical Aspects of Chemical and Biological Warfare, E. T. T. F.R. Sidell, D.R. Franz, Ed. (Office of The Surgeon General at TMM Publications, Washington, DC, 1997), vol. I, chap. 33, pp. 643–654.

16. K. H. Lewis, E. V. Hill, Practical Media and Control Measures for Producing Highly Toxic Cultures of Clostridium botulinum, Type A. J Bacteriol 53, 213–230 (1947).

17. J. T. Duff, G. G. Wright, J. Klerer, D. E. Moore, R. H. Bibler, Studies on immunity to toxins of *Clostridium botulinum*. I. A simplified procedure for isolation of type A toxin. J Bacteriol 73, 42–47 (1957).

18. H. Sugiyama, L. J. Moberg, S. L. Messer, Improved procedure for crystallization of Clostridium botulinum type A toxic complexes. Appl Environ Microbiol 33, 963–966 (1977).

19. C. J. Malizio, M. C. Goodnough, E. A. Johnson, Purification of Clostridium botulinum type A neurotoxin. Methods Mol Biol 145, 27–39 (2000).

20. M. Bradshaw, S. S. Dineen, N. D. Maks, E. A. Johnson, Regulation of neurotoxin complex expression in Clostridium botulinum strains 62A, Hall A-hyper, and NCTC 2916. Anaerobe 10, 321–333 (2004).

21. D. B. Lacy, W. H. Tepp, A. C. Cohen, B. R. DasGupta, R. C. Stevens, Crystal structure of botulinum neurotoxin type A and implications for toxicity. Nat Struct Biol 5, 1091–1104 (1996).

22. A. B. Scott, Botulinum toxin injection of eye muscles to correct strabismus. Trans Am Ophthalmol Soc 79, 734–770 (1981).

23. L. F. Lassen, J. Adams, Botulinum toxin therapy for blepharospasm in the otolaryngology clinic. ORL Head Neck Nurs 12, 12–13 (1994).

24. L. Zhang, W. J. Lin, S. Li, K. R. Aoki, Complete DNA sequences of the botulinum neurotoxin complex of Clostridium botulinum type A-Hall (Allergan) strain. Gene 315, 21–32 (2003).

25. P. K. Fang, B. H. Raphael, S. E. Maslanka, S. Cai, B. R. Singh, Analysis of genomic differences among Clostridium botulinum type A1 strains. BMC Genomics 11, 725 (2010).

26. C. H. Williamson et al., Comparative genomic analyses reveal broad diversity in botulinum-toxin-producing Clostridia. BMC Genomics 17, 180 (2016).

27. T. Kim, J. Y. Lee, S. Gil, F. Gessler, C. H. Shin, Complete genome sequence of Clostridium botulinum type A strain (X58540) isolated from cattle feces. Microbiol Resour Announc 14, e0098824 (2025).

28. M. C. Lee, C. J. Marx, Repeated, selection-driven genome reduction of accessory genes in experimental populations. PLoS Genet 8, e1002651 (2012).

29. J. P. McCutcheon, N. A. Moran, Extreme genome reduction in symbiotic bacteria. Nat Rev Microbiol 10, 13–26 (2011).

30. S. Cao, G. Brandis, D. L. Huseby, D. Hughes, Positive Selection during Niche Adaptation Results in Large-Scale and Irreversible Rearrangement of Chromosomal Gene Order in Bacteria. Mol Biol Evol 39 (2022).

31. S. Koskiniemi, S. Sun, O. G. Berg, D. I. Andersson, Selection-driven gene loss in bacteria. PLoS Genet 8, e1002787 (2012).

32. W. Sung et al., Asymmetric Context-Dependent Mutation Patterns Revealed through Mutation-Accumulation Experiments. Mol Biol Evol 32, 1672–1683 (2015).

33. H. Lee, E. Popodi, H. Tang, P. L. Foster, Rate and molecular spectrum of spontaneous mutations in the bacterium Escherichia coli as determined by whole-genome sequencing. Proc Natl Acad Sci U S A 109, E2774–2783 (2012).

34. T. T. Kibota, M. Lynch, Estimate of the genomic mutation rate deleterious to overall fitness in E. coli. Nature 381, 694–696 (1996).

35. J. E. Barrick, R. E. Lenski, Genome dynamics during experimental evolution. Nat Rev Genet 14, 827–839 (2013).

36. H. Long et al., Evolutionary determinants of genome-wide nucleotide composition. Nat Ecol Evol 2, 237–240 (2018).

37. J. H. Miller, Spontaneous mutators in bacteria: insights into pathways of mutagenesis and repair. Annu Rev Microbiol 50, 625–643 (1996).

38. A. Couce et al., Mutator genomes decay, despite sustained fitness gains, in a long-term experiment with bacteria. Proc Natl Acad Sci U S A 114, E9026–E9035 (2017).

39. V. Ng, Lin, W.J., Comparison of Transcriptomes and Sporulation of Two Clostridium botulinum A1 Strains. Adv Biotech & Micro. 11 (2018).

40. R. E. Lenski, Experimental evolution and the dynamics of adaptation and genome evolution in microbial populations. ISME J 11, 2181–2194 (2017).

41. T. F. Cooper, R. E. Lenski, Experimental evolution with E. coli in diverse resource environments. I. Fluctuating environments promote divergence of replicate populations. BMC Evol Biol 10, 11 (2010).

42. N. A. Moran, Microbial minimalism: genome reduction in bacterial pathogens. Cell 108, 583–586 (2002).

43. C. Toft, S. G. Andersson, Evolutionary microbial genomics: insights into bacterial host adaptation. Nat Rev Genet 11, 465–475 (2010).

44. C. H. Kuo, H. Ochman, Deletional bias across the three domains of life. Genome Biol Evol 1, 145–152 (2009).

45. R. Albalat, C. Canestro, Evolution by gene loss. Nat Rev Genet 17, 379–391 (2016).

46. O. Tenaillon et al., Tempo and mode of genome evolution in a 50,000-generation experiment. Nature 536, 165–170 (2016).

47. R. E. Lenski, M. Travisano, Dynamics of adaptation and diversification: a 10,000-generation experiment with bacterial populations. Proc Natl Acad Sci U S A 91, 6808–6814 (1994).

48. S. Wielgoss et al., Mutation rate dynamics in a bacterial population reflect tension between adaptation and genetic load. Proc Natl Acad Sci U S A 110, 222–227 (2013).

49. H. J. Muller, The Relation of Recombination to Mutational Advance. Mutat Res 106, 2–9 (1964).

50. P. Funchain et al., The consequences of growth of a mutator strain of Escherichia coli as measured by loss of function among multiple gene targets and loss of fitness. Genetics 154, 959–970 (2000).

51. G. D’Souza et al., Less is more: selective advantages can explain the prevalent loss of biosynthetic genes in bacteria. Evolution 68, 2559–2570 (2014).

52. E. R. Moxon, P. B. Rainey, M. A. Nowak, R. E. Lenski, Adaptive evolution of highly mutable loci in pathogenic bacteria. Curr Biol 4, 24–33 (1994).

53. T. Swings et al., Adaptive tuning of mutation rates allows fast response to lethal stress in Escherichia coli. Elife 6 (2017).

54. F. J. A. Lebeda, M.; Dembek, Z.F, Yesterday and Today: The Impact of Research Conducted at Camp Detrick on Botulinum Toxin. Military Medicine 183, 11 (2018).

55. F. N. Fu, S. K. Sharma, B. R. Singh, A protease-resistant novel hemagglutinin purified from type A Clostridium botulinum. J Protein Chem 17, 53–60 (1998).

56. T. J. Smith et al., Analysis of the neurotoxin complex genes in Clostridium botulinum A1-A4 and B1 strains: BoNT/A3, /Ba4 and /B1 clusters are located within plasmids. PLoS One 2, e1271 (2007).

57. M. Martin, Cutadapt removes adapter sequences from high-throughput sequencing reads. EMBnet 17 (2011).

58. A. Bankevich et al., SPAdes: a new genome assembly algorithm and its applications to single-cell sequencing. J Comput Biol 19, 455–477 (2012).

59. B. D. Ondov et al., Mash: fast genome and metagenome distance estimation using MinHash. Genome Biol 17, 132 (2016).

60. J. Felsenstein, Mathematics vs. Evolution: Mathematical Evolutionary Theory. Science 246, 941–942 (1989).

61. N. Saitou, M. Nei, The neighbor-joining method: a new method for reconstructing phylogenetic trees. Mol Biol Evol 4, 406–425 (1987).

62. A. E. Darling, B. Mau, N. T. Perna, progressiveMauve: multiple genome alignment with gene gain, loss and rearrangement. PLoS One 5, e11147 (2010).

63. S. Kurtz et al., Versatile and open software for comparing large genomes. Genome Biol 5, R12 (2004).

64. H. Li, Minimap2: pairwise alignment for nucleotide sequences. Bioinformatics 34, 3094–3100 (2018).

65. M. A. DePristo et al., A framework for variation discovery and genotyping using next-generation DNA sequencing data. Nat Genet 43, 491–498 (2011).

66. A. McKenna et al., The Genome Analysis Toolkit: a MapReduce framework for analyzing next-generation DNA sequencing data. Genome Res 20, 1297–1303 (2010).

67. W. Huang, L. Li, J. R. Myers, G. T. Marth, ART: a next-generation sequencing read simulator. Bioinformatics 28, 593–594 (2012).

68. A. L. Delcher, A. Phillippy, J. Carlton, S. L. Salzberg, Fast algorithms for large-scale genome alignment and comparison. Nucleic Acids Res 30, 2478–2483 (2002).

69. J. W. Sahl et al., NASP: an accurate, rapid method for the identification of SNPs in WGS datasets that supports flexible input and output formats. Microb Genom 2, e000074 (2016).

70. P. Cingolani et al., A program for annotating and predicting the effects of single nucleotide polymorphisms, SnpEff: SNPs in the genome of Drosophila melanogaster strain w1118; iso-2; iso-3. Fly (Austin) 6, 80–92 (2012).

71. B. Q. Minh et al., IQ-TREE 2: New Models and Efficient Methods for Phylogenetic Inference in the Genomic Era. Mol Biol Evol 37, 1530–1534 (2020).

72. S. Kalyaanamoorthy, B. Q. Minh, T. K. F. Wong, A. von Haeseler, L. S. Jermiin, ModelFinder: fast model selection for accurate phylogenetic estimates. Nat Methods 14, 587–589 (2017).

73. K. P. Schliep, phangorn: phylogenetic analysis in R. Bioinformatics 27, 592–593 (2011).

## References

1. Williamson, C.H. et al. Comparative genomic analyses reveal broad diversity in botulinum-toxin-producing Clostridia. BMC Genomics 17, 180 (2016).

2. Smith, T., Williamson, C., Hill, K., Sahl, J. & Keim, P. Botulinum neurotoxin-producing bacteria. Isn’t it time that we called a species a species? mBio 9:e01469–18., supplementary information (2018).

3. Fang, P.K., Raphael, B.H., Maslanka, S.E., Cai, S. & Singh, B.R. Analysis of genomic differences among Clostridium botulinum type A1 strains. BMC Genomics 11, 725 (2010).

4. Kurtz, S. et al. Versatile and open software for comparing large genomes. Genome Biol 5, R12 (2004).

